# Impacts of Deforestation and Land Use/Land Cover Change on Carbon Stock and Dynamics in the Jomoro District, Ghana

**DOI:** 10.1101/2024.04.26.591345

**Authors:** Elisa Grieco, Elia Vangi, Tommaso Chiti, Alessio Collalti

## Abstract

Tropical deforestation in the African continent plays a key role in the global carbon cycle and bears significant implications in terms of climate change and sustainable development. Especially in Sub-Saharan Africa, where more than two-thirds of the population rely on forest and woodland resources for their livelihoods, deforestation and land use changes for crop production lead to a substantial loss of ecosystem-level carbon stock. Unfortunately, the impacts of deforestation and land use change can be more critical than in any other region, but these are poorly quantified. We analyse changes in the main carbon pools (above- and below-ground, soil and litter, respectively) after deforestation and land use/land cover change, for the Jomoro District (Ghana), by assessing the initial reference level of carbon stock for primary forest and the subsequent stock changes and dynamics as a consequence of conversion to the secondary forest and to six different tree plantations (rubber, coconut, cocoa, oil palm, and mixed plantations). Results indicate overall a statistically significant carbon loss across all the land uses/covers and for all the carbon pools compared to the primary forest with the total carbon stock loss ranging between 85% and 35% but with no statistically significant differences observed in the comparison between primary forest and mixed plantations and secondary forest. Results also suggest that above-ground carbon and soil organic carbon are the primary pools contributing to the total carbon stocks but with opposite trends of carbon loss and accumulation. Strategies for sustainable development, policies to reduce emissions from deforestation and forest degradation, carbon stock enhancement (REDD+), and planning for sustainable land use management should carefully consider the type of conversion and carbon stock dynamics behind land use change for a win-win strategy while preserving carbon stocks potential in tropical ecosystems.

## 1 Introduction

The increasing decline of African forests is the result of land use change related mostly to anthropogenic activities, affecting not only biodiversity, carbon (C) and water cycle but also undermining the mitigation potential of tropical ecosystems. A relevant aspect of climate change mitigation strategies is the understanding of the dynamics of land use changes following deforestation (Masolele et al., 2024). Despite the widely recognised importance of the Sub-Saharan Africa region to the global C-budget, the knowledge gap at C-pool level still needs to be filled (Houghton & Hackler, 2006; Olorunfemi et al., 2022). Land use change (LUC) activities related to deforestation play a major role in determining sources and sinks of carbon (Le Quéré et al., 2009) being responsible for emissions of about 1.9 PgC yr^−1^ over the period 2013-2022 (Friedlingstein et al., 2023). Africa experienced the largest annual rate with 3.9 million hectares of forests lost between 2010 and 2020 with an increasing trend since 1990 and long-term management plans existing for less than 25% of forests, one of the lowest levels after South America (20%)(FAO, 2020). Agricultural expansion has a key role in the African deforestation process, leading to 64% of total forest loss, in favour of small-scale cropland, from 2001 to 2020. Moreover, 7% of forest conversion in Africa has been attributed to commodity crops such as rubber, oil palm and cocoa, common in West Africa, notably in Ghana (Masolele et al., 2024). An IUCN analysis (IUCN, 2008) highlighted the critical situation of Ghana’s Western Region where most of the patches of forestland outside forest reserves, that existed in 1990, had been converted to other land uses by 2007. Findings from Ampim et al. (2021) also confirm a loss in forest cover in the Western Region (∼704.7 km^2^) between 1995 and 2019 despite a general increase at the national level (+23.3%). National policies, aiming at modernising and expanding tree crop plantation areas while supporting smallholder farmers, are also influenced by global demand in the supply chain and the need to boost employment and local food security (Ampim *ibid.*). Tree plantations could apparently seem a perfect win-win solution for enhancing local community livelihood while promoting climate change mitigation and economic development, probably even more than pure afforestation and reforestation practices (Kongsager et al., 2013). Nevertheless, it is questionable whether this choice can be sustainable in terms of long-term C-stock balance at the ecosystem level. As also underlined by Pendrill et al. (2022), land-use change research in Africa and ‘commodity-specific land use dynamics data’ are poorly known and efforts should be focused on better characterising deforestation in smallholder shifting agriculture, being one of the main drivers of deforestation. At the same time, the expansion of commodity crops like cocoa, rubber and oil palm in large-scale cropland in humid and dry forests in western and south-eastern Africa, indicates a likely vulnerability to future land use change and a real obstacle to zero deforestation supply chain (Masolele et al., 2024). Information on C-sequestration potentialities and C-stock dynamics of tree crop monoculture systems in developing countries – and particularly in Africa – are scarce and incomplete (Kongsager *ibid.*). It is undeniable that ecosystems’ resilience, forest resource protection, and mitigation potentialities are interlinked with socioeconomic improvements but more research is needed to properly assess the actual impact of forest reduction for tree crop expansion on tropical ecosystems. This study aims to untangle some unclear assumptions on the supposed beneficial effects of tree crop plantations in terms of C-stock potentials. More specifically, the present work aims to address the following questions by analysing data and describing and discussing results from the Jomoro District in Ghana (Africa): 1) What are the effects of land use/land cover change in tropical forested land, on different carbon pools compared to the primary forest?; 2) What are the changes in relation to the establishment of different tree plantations?; 3) What is the range, rate and magnitude of these variations and the dynamics in terms of C-stocks across different C-pools and does it depend on the considered pool, the age or type of conversion?

## 2. Material and methods

### 2.1. Study area

The Jomoro District occupies the Southwestern corner of the Western Region of Ghana and covers an area of 1.344 km^2^, about 5.6% of the total area of the Western Region (JDA, 2009, 2014). The district has extensive rainforest and the south-central part includes the Ankasa Forest Reserve. The potential vegetation is represented by high-forests, not uniform throughout but divided into several belts which differ in their floristic composition, general characters and distribution mainly related to rainfall and soil acidity. The high forest is characterised by species referred to as indicator trees, such as *Cynometra ananta* Hutch. & Dalziel, *Lophira alata* Banks ex Gaertn. and *Tarrietia utilis* Sprague also known as the *Cynometra-Lophira-Tarrietia* association (Ahn, 1961). The district lies on four main geological formations: Granites, Lower Birrimian, the Tertiary Sands and the Coastal Sands (Hall & Swaine, 1976; Schlüter, 2005; Chiti et al., 2013). The soils developed from the weathering of granite (Atsivor et al., 2001) can be divided into two main groups in the Jomoro District: Ochrosols and Oxysols (Soil Taxonomy, 2010). The two groups differ mainly in the topsoil with the Ochrosols showing a pH of 5.5 and the Oxisols of less than 5 (Ahn, 1961). The landscape is characterised by slopes varying from 1 to 30% (Wauters et al., 2008). The climate is classified as Equatorial Monsoon (Kottek et al., 2006). The rainfall regime in the region is bimodal with four seasons; two rainy seasons: a major in April-July and a minor in October-November, and two dry seasons: a major in December-March and a minor in August-September. Monthly rainfall varies between 0 and 500 mm, while annual rainfall ranges from 1200 to 1800 mm. The average relative air humidity is between 95 and 100%. The average annual temperature ranges between 24 and 27 °C. Absolute extreme temperatures are 15 and 40 °C. The main drivers of deforestation and forest degradation in the high forest zone of Ghana are strictly braided into multiple factors which encompass agricultural expansion, mining, wood/timber extraction and infrastructure extension (Hansen et al., 2009; Ampim et al., 2021). Among the dominant drivers of forest loss for agricultural expansion, is the increased expansion of cocoa, oil palm and rubber with a forest conversion of 25.4, 1.9 and 1.5%, respectively, with 25.9% of forest converted into small-scale cropland between 2001 and 2020 (Masolele et al., 2024).

### 2.2. Data collection

#### 2.2.1. Sites’ selection

The sites’ selection aimed at locating the main land uses after the original rainforest clearance throughout the Jomoro District and to this end, selection was based on results from interviews with local farmers and on field surveys. It was considered ‘eligible’, the site which has been deforested to be used for only a single plantation type ever since. The eligibility criteria have been applied to assess the changes in Total Carbon Stocks (TCS) as a direct effect of the forest clearing on the main C-pools: above-ground biomass, below-ground biomass, soil, litter, and standing dead wood. The sampling fieldwork was carried out on each site, in three selected plots per site for soil, litter and above-ground then followed by laboratory and data analysis. Extensive surveys, performed in 2009 and 2010 within the Jomoro District, have allowed the identification, in addition to forest and secondary forest sites, of five main commodities plantations for a total of 23 sites (i.e. 3 for forest, 2 for secondary forest, 2 for oil palm, 2 for cocoa, 4 for rubber, 2 for mixed and 8 for coconut plantation)(Figure 1, Table S1 in Supplementary Material).

**Figure 1.**
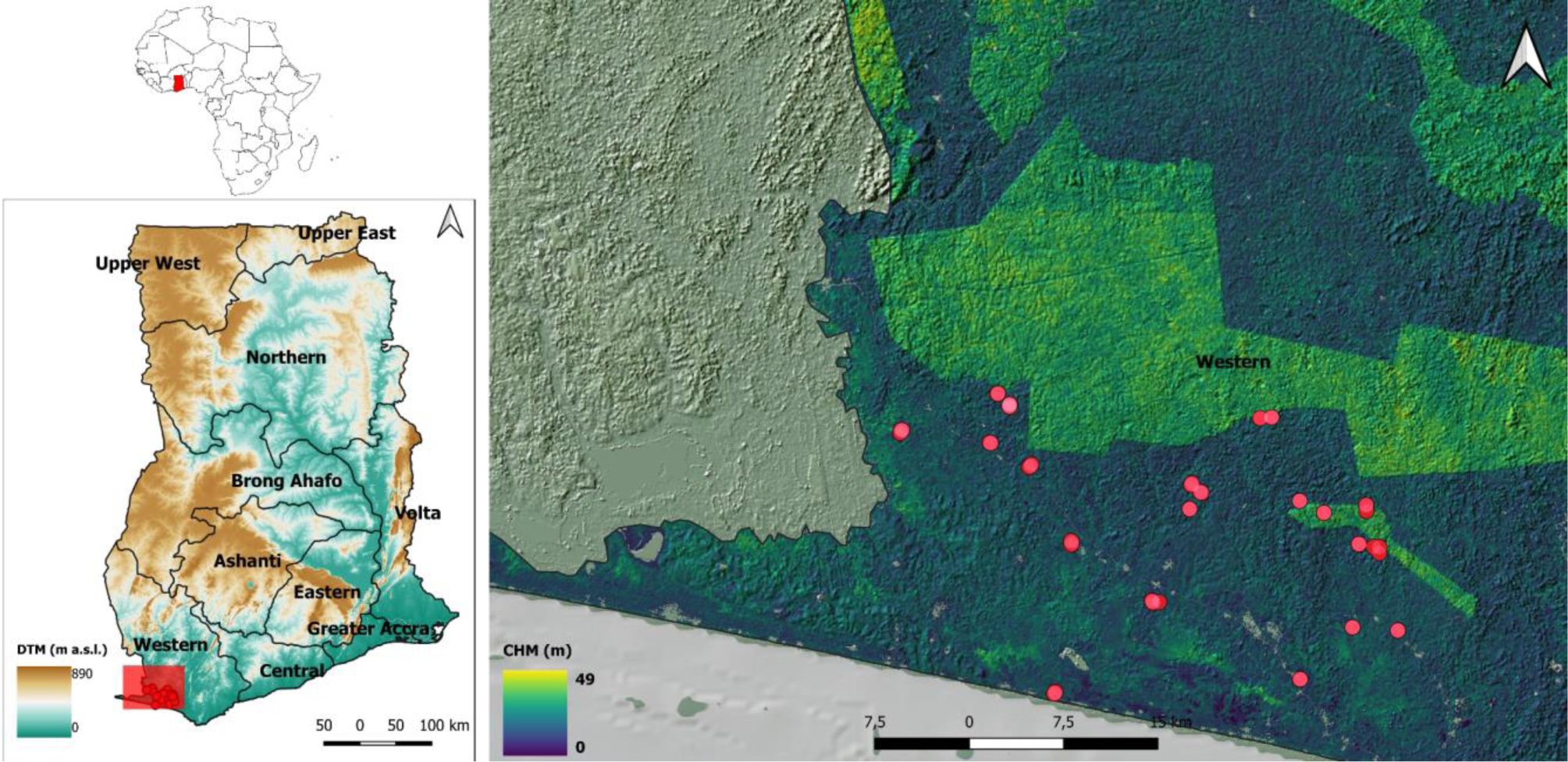
Map of the survey area and the selected sites (red dots) in the Jomoro District, Western Region, Ghana.

##### Primary forests

Two primary forest sites were identified. The former was a patch of land close to Cocotown village, preserved as a high-forest due to its sacred value for local people. The utilisation of wood and collections of other woody products was strictly forbidden. The second site was a portion of the Ankasa conservation area. The sites were identified with the codes ‘ANKF’ and ‘CTF’, respectively. A third patch of primary forest was selected near Bawia village. This parcel of land was designated as community property, reserved for future farming activities.

##### Secondary forests

The secondary forest sites were two, ‘SF_10_’ and ‘SF_20_’. The first was deforested in 2000 and then abandoned. Similarly, on the second site, the pristine forest was cleared in 1988, and cultivated for one year with cassava and coconut but farming activities did not succeed because of pests, so the land was then abandoned.

##### Oil palm plantations

Two oil palm plantation sites were identified, having been established 8 and 25 years before the survey. These plantations were established immediately after clearing the primary forest originally covering those areas. The plantation ‘OP_8_’ was established in 2002 while ‘OP_25_’ was established in 1985, one year after forest clearance. At the time of the survey, the plantation OP_25_ was in its second generation. The replacement was done in 2005, so that, at the time of the survey, the new plantation was only 4-year-old.

##### Cocoa plantations

Two cocoa plantations were identified, the first at Cocotown village, along the river Tano. This area has been historically famous for cocoa production since the 18th century. The chosen site, ‘CC_120_’, was established in 1890 and continuously cultivated with cocoa. The assessed plantation was reported never to have been replaced because the trees were left for natural regeneration through coppicing. The second site, ‘CC_34_’, was selected in a village located south of Cocotown. The plantation was established in 1975 following the clearance of the primary forest and has been consistently cultivated cocoa ever since.

##### Coconut plantations

Height sites for coconut plantations were identified. The first, ‘CN_21_’, was located near the western entrance of the Ankasa conservation area and was established in 1988. The second site, ‘CN_28_’, was situated at Nyamenle Keya Beven near Sowodzadzem village. It was established in 1982, while the third site ‘CN_44_’ was located between Navrongo crossroad and Abudu village and was established in 1966. The fourth, ‘CN_15_’ was located close to Betekomoa village and was established in 1995, the fifth, ‘CN_50_’, was near Aboyele village and was established in 1960. The sixth one, ‘CN_100_’, was close Nuba village and was established in 1910 and, at the time of the survey, the plantation was 100-year-old. The seventh site, ‘CN_53_’, was deforested in 1956 while the eighth, ‘CN_95_’, was in 1915.

##### Rubber plantations

Two sites deforested at different times were found also for rubber. The first site,‘RP_5_’, is located near Johnatan village and was established in 2005. The second rubber plantation,‘RP_10_’, located near New Ankasa village, was established in 2000. At both sites were present the first generation of trees. The rubber plantations found on this substrate were two, the first, ‘RP_14_’, located near Bawia village was established in 1996, while the second one, ‘RP_50_’, located in Mpataba village was established in 1960.

##### Mixed plantations

The investigated sites cultivated as mixed plantations in the Jomoro district were two. The tree component has been considered for measurements. The first plantation, ‘MP_36_’, was established in 1974, near Agaege village. Trees proportion within the plantation was oil palm (40%) and coconut (60%). The latter site, ‘MP_50_’, was established in 1960, in the proximity of one of the Ankasa Conservation area. This mixed plantation was characterised by a composition of oil palm (38%), coconut (19%) and other species (13%).

#### 2.2.2. Sampling activity and data analysis

##### Above- and below-ground biomass

To assess the above-ground biomass (AGB), we conducted morphometric measurements of the standing vegetation starting by measuring tree height (H, in meters) and diameter at breast height (DBH, in cm), according to FAO protocol (FAO; Ponce-Hernandez, 2004). The DBH measurement threshold was 5 cm according to VCS (VCS, 2010). The sample plots’ dimensions were 10 by 10 meters, coinciding with the soil sample plots. For species with a wider tree planting spacing, the plot dimensions were doubled (e.g. coconut). The tree height has been measured with a Suunto clinometer, the DBH with a dendrometer calliper, and the sampling area bordered with a measuring tape. Each plot within the selected site has been georeferenced through a GPS device (Garmin/GPS60). Dead-standing trees were also considered in measurements and included in the deadwood biomass pool. Non-destructive method was adopted for AGB estimation, through the use of allometric equations (Table 1). The below-ground biomass (BGB) was not collected on the field directly but estimated as a percentage of the above-ground (AGB) biomass using the root-to-shoot ratio as in Cairns et al. (1997), Houghton et al. (2001), Achard et al. (2002), Ponce-Hernandez, (2004), Mokany et al. (2006) and Ramankutty et al. (2007). BGB has been considered 20% of the AGB based on a predictive relationship established from extensive literature reviews (Cairns et al., 1997; Houghton et al., 2001; Achard et al., 2002; Ponce-Hernandez, 2004; Mokany et al., 2006; Ramankutty et al., 2007).

**Table 1.**
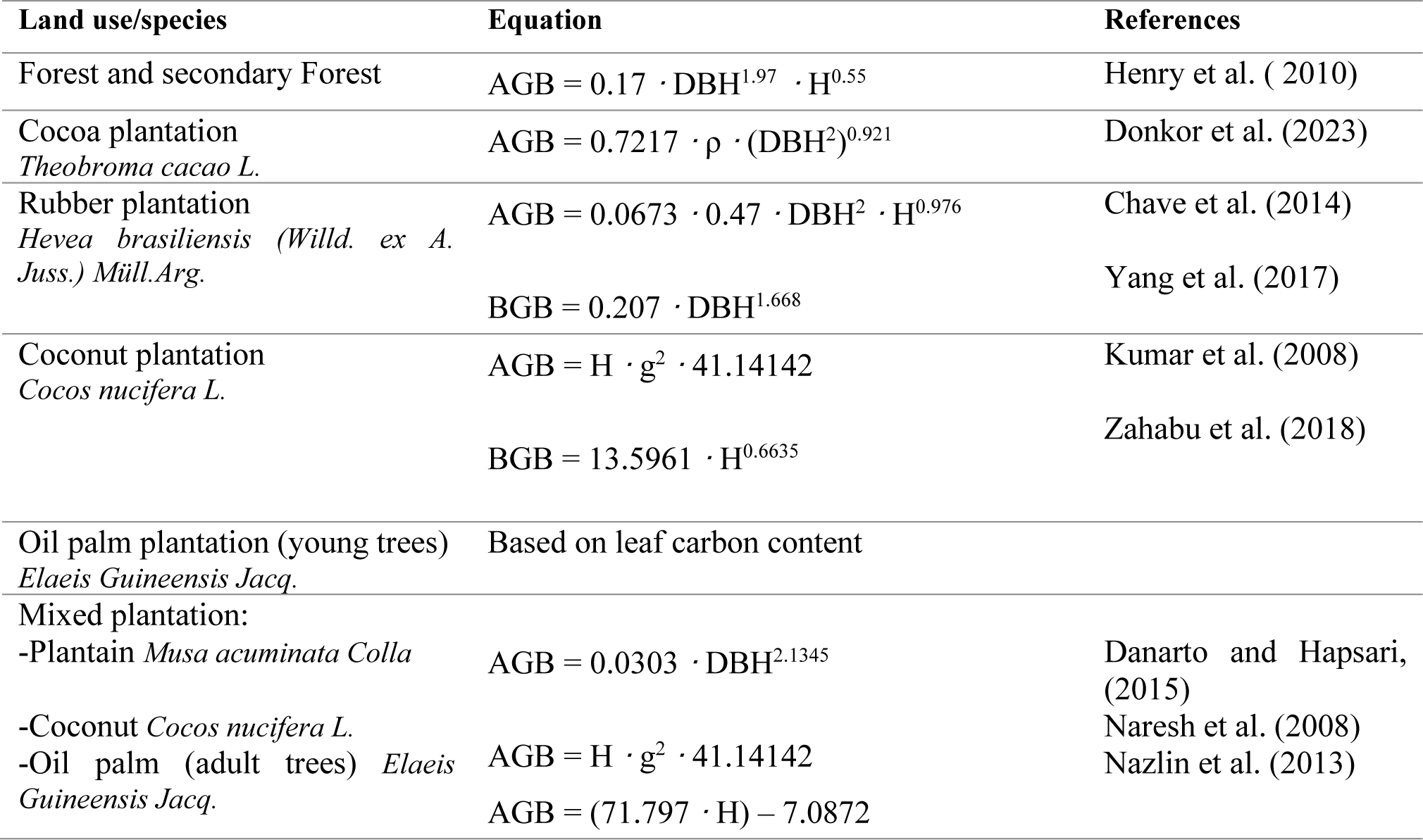
Allometric equations used to assess Above-Ground Biomass (AGB) and Below-Ground Biomass (BGB) in this study. AGB and BGB are expressed in kg of Dry Matter per tree, Diameter at Breast Height (DBH) in cm, Tree Height (H) in m, wood density (ρ) in g·cm^−3^ and girth (g) in m.

##### Soil and Litter

For assessing the changes in soil organic carbon stock, it was adopted the soil sampling protocol proposed by Stolbovoy et al. (2005) and Chiti et al. (2014). Three random sampling plots for each land-use system have been considered. In each plot, 20 by 20 m, soil samples were collected along a grid of 25 sampling points at three depths: 0-10, 10-20, 20-30 cm. Then the 25 samples for each depth were pooled together to have three composite samples per depth and per land use area. In the middle of each plot a minipit was opened to collect bulk density samples at the same depths, to have one sample per depth per plot and three samples per depth and per land use area, following the core method (Blake and Hartge, 1986). Soil samples were oven-dried at 60 °C, and sieved at 2 mm to separate the rock fragments from the fine earth. Both of the fractions were weighed. Fine earth was analyzed for total carbon and nitrogen (N) using CN analyzer by dry combustion (Thermo Finnigan Flash EA112 CHN). Bulk density samples were oven-dried at 105 °C until constant mass, then the oven-dry weights of the soil samples were divided by the cylinder volume and calculated as Mg m^−3^. The soil organic carbon stock was calculated as follows:

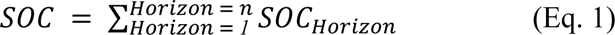

Where

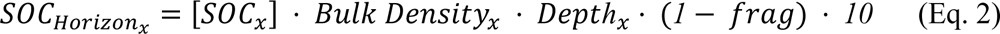

where SOC is soil C-content per unit area (MgC ha^−1^), [SOC] is carbon concentration in soil sample (kgC kg^−1^ soil), the *Bulk Density* is the soil density of the fine earth expressed as (Mg m^−3^), *Depth* is the thickness of the horizon within the considered section (cm) and *frag* is the percentage of rock fragments, and *x* is for the different soil horizons (Poeplau et al. 2017).

Where present, the litter layer was collected within a frame 40 by 40 cm (e.g., in the case of primary and secondary forests, rubber and cocoa plantations) and 3 by 3 m in case of big leaves such as those of the oil palm plantations. In the rest of the cases, the number of collected samples was 5 in each plot for a total amount of 15 in each site. For the species with big leaves, a single leaf was collected in each plot and it has been counted as the number of the leaves in the 3 by 3 m frame. The litter samples were oven-dried at 60 °C, weighed, grounded and analysed for total C and N by dry combustion (Thermo Finnigan Flash EA112 CHN). The C-stock was estimated by multiplying the weight of dry-matter of the sample by the C-concentration and reporting the value on a surface basis, as follows:

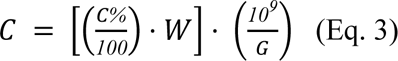

Where C is the final value of C in the litter layer (MgC ha^−1^), C% is the percentage of carbon concentration in the considered sample, W is the weight of the litter sample and G is the surface of the grid (cm^3^).

##### Deadwood biomass

Dead wood biomass was estimated in terms of dead-standing trees, expressed as a percentage of the AGB.

##### From biomass to carbon

According to IPCC (2003), Losi et al. (2003), Sarmiento et al. (2005), Chave et al. (2005), Pearson et al. (2005), and Fonseca et al. (2012), a coefficient of 0.5 was used to convert tree biomass to carbon. Carbon per tree, expressed in kg, was then multiplied by the number of trees per hectare to determine C in above- and below-ground biomass on an area basis (MgC ha^−1^). The exception was made for *Musa acuminata*, for which a conversion factor of 0.46 was used (Hairiah et al., 2010; Danatro and Hapsari, 2015). Overall the analyses, including the statistical ones, were performed, described and discussed at the carbon level.

### 2.3 Statistical Analysis

Statistical tests were performed to check for significant differences in C-pools across different land uses/covers. We fit a one-factor ANOVA to test if land uses have an overall significant effect, and if so, we tested all pairwise comparisons between the seven land uses. For this purpose, the post hoc general linear hypothesis test (glht)(p =0.05) was used to test multiple hypotheses concerning a linear function of interest obtained from the matrix of parametric contrasts calculated on the land use factor by Tukey’s method (Bretz et al., 2010). We also performed a Dunnett post hoc test (p =0.05) using the forest land use as a control group. We adjusted the p-value based on the joint normal distribution of the linear function (Dunnett & Tamhane, 1991). We used a sandwich estimator that provides a heteroscedasticity-consistent covariance matrix estimate for both post hoc tests when needed. Before proceeding, normality and homogeneity of variance were checked via the Shapiro-Wilk and Levene’s tests, respectively (for both p =0.05). All analyses were performed using the statistical software R, with the package *‘multcomp’* and *‘sandwich’* (Zeileis, 2006).

## 3. Results

Results for all carbon pools (as singularly taken and overall accounted in TCS) are described in comparison to forest and other land uses/covers and in absolute terms as also considering differences, within the same land use/cover, between ages.

### 3.2 Carbon stocks and dynamics

#### Above-ground carbon (AGC)

The overall differences in AGC between forest, secondary forest and all the plantation types resulted to be statistically significant (p <0.05) except for mixed plantation (p =0.19) (Fig.2). Compared to the pristine forest, all the plantations show a diminished AGC which varies both with the age and the type of plantation (Table 2). It is noteworthy that, considering the age of the plantations, the AGC of secondary forest (SF_10_), rubber (RP_14_ and RP_50_), mixed plantation (MP_36_, MP_50_) and coconut (CN_95_), is not statistically different from the forest (Figure S1 in Supplementary Material). The AGC from seven subplots of primary forest, results to be, on average, 263.9 ±8.5 MgC ha^−1^ (here and elsewhere, ‘±’ denotes one standard deviation). This was considered the reference value to be confronted with. Secondary forest resulted in an AGC of 78.7 ±5.2 (SF_10_) and 33.1 ±2.5 MgC ha^−1^ (SF_20_), showing a loss of 70.2 and 87.5%, respectively (Table 3). After deforestation, the observed mean AGC annual increment was 7.9 in SF_10_ and 1.65 MgC ha^−1^ yr^−1^ in SF_20_. In oil palm plantations, the AGC accounts for 3.8 ±0.03 (4-year-old) and 3.4 ±0.1 MgC ha^−1^ (8-year-old) with a statistically significant loss from the forest reference level of 98.6 and 98.7%, respectively. The estimated mean AGC annual increment was 1 ±0.5 and 0.4 ±0.1 MgC ha^−1^ yr^−1^ in 4 and 8-year-old stands respectively. For the 120-year-old cocoa plantation, AGC resulted in 19.5 ±0.1 while the 34-year-old stand has 21.2 ±0.3 MgC ha^−1^. The loss in this case accounts respectively for 92.6 and 92% with a mean AGC annual increment since deforestation, accounting for 0.6 ±0.2 and 0.2 ±0.05 MgC ha^−1^ yr^−1^ in CC_34_ and CC_120_. Four rubber plantation AGC stock ranges between 10.7 ±0.3 (RP_5_) and 162.9 ±5 MgC ha^−1^ (RP_50_). The C-loss in this case varies from 95.9 to 38.3%, respectively, while the mean AGC annual increment since forest clearing changes from 2.1 ±0.4 to 3.3 ±0.7 MgC ha^−1^ yr^−1^, in RP_5_ and RP_50_ respectively. The AGC in MP_36_ was 109.02 ±3.73 and 164.22 ±8.20 MgC ha^−1^ in MP_50_, with a loss accounting for 58.7 and 37.8%, respectively. In terms of mean annual AGC increment, MP_36_ was 3 ±0.5 while MP_50_ was 3.3 ±2.1 MgC ha^−1^ yr^−1^. Coconut AGC ranges from 44.4 ±0.5 (CN_15_), to 46.7 ±1.7 MgC ha^−1^ (CN_100_). The AGC loss on the eight coconut sites varies between 93.3 (CN_21_) to 56.7% (CN_95_) while the mean annual AGC increment changes from 3 ±0.08 in CN_15_ to 0.5 ±0.2 MgC ha^−1^ yr^−1^ in CN_100_ (Table 2 and 3, and Table S2 in Supplementary Material).

**Figure 2.**
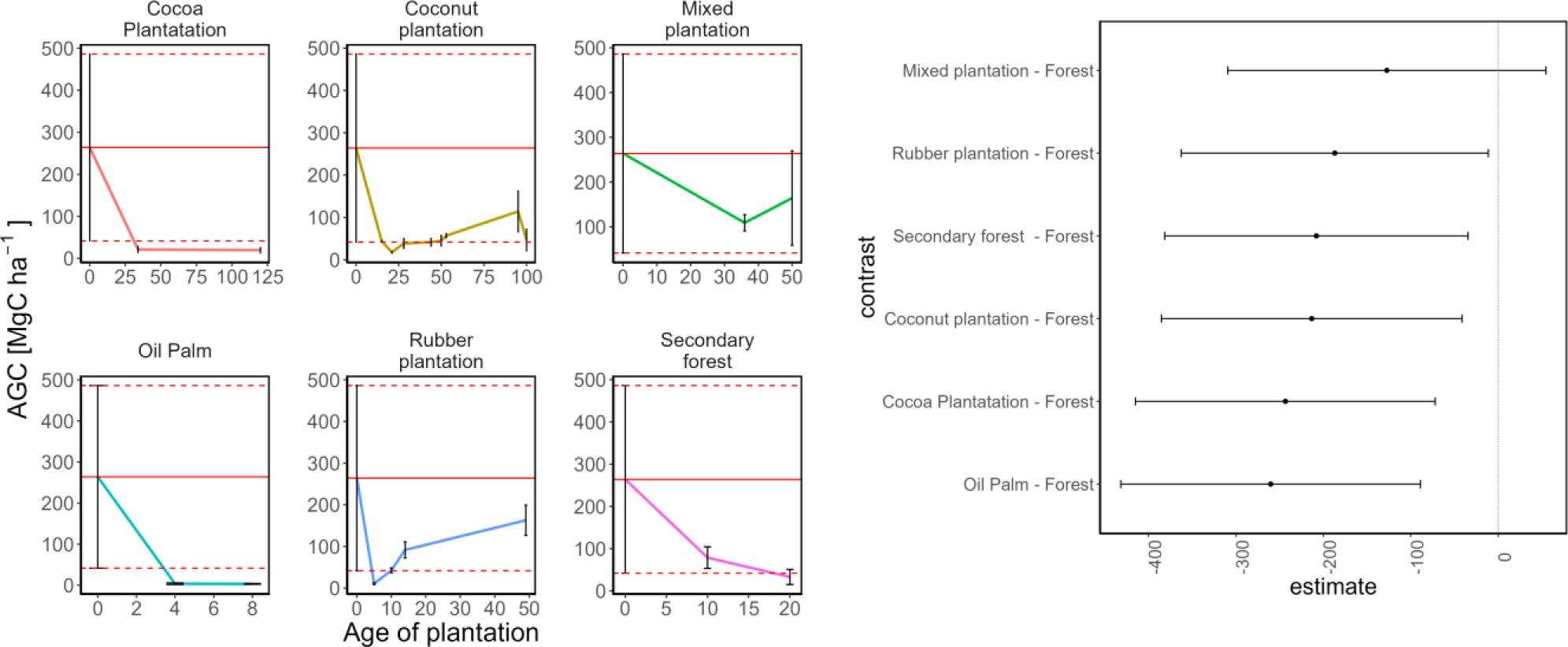
Above-Ground Carbon (AGC in MgC ha^−1^) stocks dynamics in secondary forest and tree plantations (left panel). Statistical significance is represented with Dunnet plot, where confidence interval crossing reference dotted line (0) indicates no statistical difference (p > 0.05)(right panel).

**Table 2.**
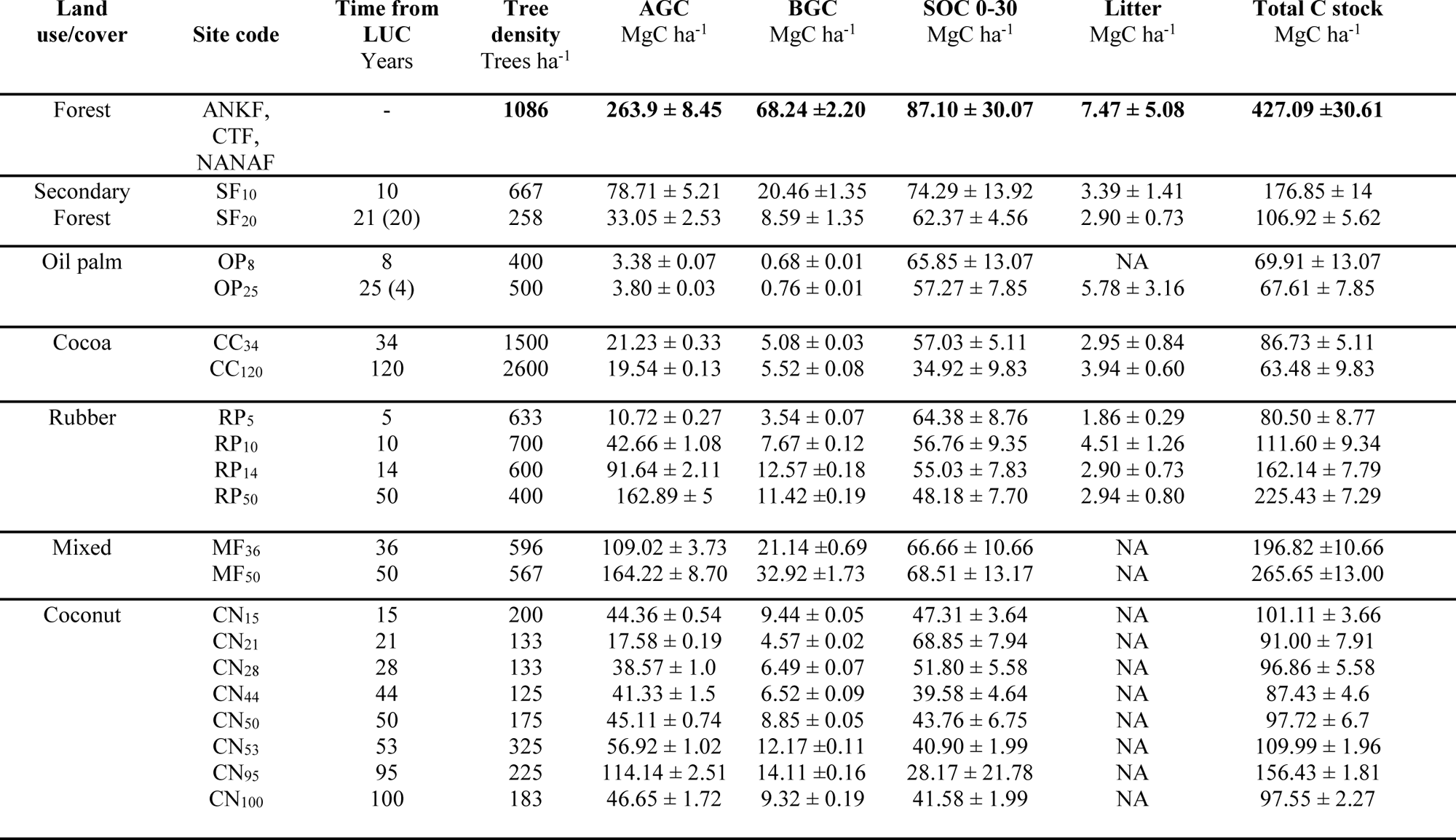
Carbon stocks of different pools (AGC, BGC, SOC, Litter) and TCS (AGC + BGC + Litter + SOC) in the forest, secondary forest and tree plantations (in MgC ha^−1^). The subscript number on the site code indicates the years since deforestation, in parentheses the age of the plantation, ‘±’ denotes one standard deviation. NA = Not Available Data.

| Land use/cover | Site code | Time from LUC Years | Tree density Trees ha<sup>-1</sup> | AGC MgC ha<sup>-1</sup> | BGC MgC ha<sup>-1</sup> | SOC 0-30 MgC ha<sup>-1</sup> | Litter MgC ha<sup>-1</sup> | Total C stock MgC ha<sup>-1</sup> |
| --- | --- | --- | --- | --- | --- | --- | --- | --- |
| Forest | ANKF, CTF, NANAF | - | 1086 | 263.9 ± 8.45 | 68.24 ± 2.20 | 87.10 ± 30.07 | 7.47 ± 5.08 | 427.09 ± 30.61 |
| Secondary Forest | SF<sub>10</sub> | 10 | 667 | 78.71 ± 5.21 | 20.46 ± 1.35 | 74.29 ± 13.92 | 3.39 ± 1.41 | 176.85 ± 14 |
|  | SF<sub>20</sub> | 21 (20) | 258 | 33.05 ± 2.53 | 8.59 ± 1.35 | 62.37 ± 4.56 | 2.90 ± 0.73 | 106.92 ± 5.62 |
| Oil palm | OP<sub>8</sub> | 8 | 400 | 3.38 ± 0.07 | 0.68 ± 0.01 | 65.85 ± 13.07 | NA | 69.91 ± 13.07 |
|  | OP<sub>25</sub> | 25 (4) | 500 | 3.80 ± 0.03 | 0.76 ± 0.01 | 57.27 ± 7.85 | 5.78 ± 3.16 | 67.61 ± 7.85 |
| Cocoa | CC<sub>34</sub> | 34 | 1500 | 21.23 ± 0.33 | 5.08 ± 0.03 | 57.03 ± 5.11 | 2.95 ± 0.84 | 86.73 ± 5.11 |
|  | CC<sub>120</sub> | 120 | 2600 | 19.54 ± 0.13 | 5.52 ± 0.08 | 34.92 ± 9.83 | 3.94 ± 0.60 | 63.48 ± 9.83 |
| Rubber | RP<sub>5</sub> | 5 | 633 | 10.72 ± 0.27 | 3.54 ± 0.07 | 64.38 ± 8.76 | 1.86 ± 0.29 | 80.50 ± 8.77 |
|  | RP<sub>10</sub> | 10 | 700 | 42.66 ± 1.08 | 7.67 ± 0.12 | 56.76 ± 9.35 | 4.51 ± 1.26 | 111.60 ± 9.34 |
|  | RP<sub>14</sub> | 14 | 600 | 91.64 ± 2.11 | 12.57 ± 0.18 | 55.03 ± 7.83 | 2.90 ± 0.73 | 162.14 ± 7.79 |
|  | RP<sub>50</sub> | 50 | 400 | 162.89 ± 5 | 11.42 ± 0.19 | 48.18 ± 7.70 | 2.94 ± 0.80 | 225.43 ± 7.29 |
| Mixed | MF<sub>36</sub> | 36 | 596 | 109.02 ± 3.73 | 21.14 ± 0.69 | 66.66 ± 10.66 | NA | 196.82 ± 10.66 |
|  | MF<sub>50</sub> | 50 | 567 | 164.22 ± 8.70 | 32.92 ± 1.73 | 68.51 ± 13.17 | NA | 265.65 ± 13.00 |
| Coconut | CN<sub>15</sub> | 15 | 200 | 44.36 ± 0.54 | 9.44 ± 0.05 | 47.31 ± 3.64 | NA | 101.11 ± 3.66 |
|  | CN<sub>21</sub> | 21 | 133 | 17.58 ± 0.19 | 4.57 ± 0.02 | 68.85 ± 7.94 | NA | 91.00 ± 7.91 |
|  | CN<sub>28</sub> | 28 | 133 | 38.57 ± 1.0 | 6.49 ± 0.07 | 51.80 ± 5.58 | NA | 96.86 ± 5.58 |
|  | CN<sub>44</sub> | 44 | 125 | 41.33 ± 1.5 | 6.52 ± 0.09 | 39.58 ± 4.64 | NA | 87.43 ± 4.6 |
|  | CN<sub>50</sub> | 50 | 175 | 45.11 ± 0.74 | 8.85 ± 0.05 | 43.76 ± 6.75 | NA | 97.72 ± 6.7 |
|  | CN<sub>53</sub> | 53 | 325 | 56.92 ± 1.02 | 12.17 ± 0.11 | 40.90 ± 1.99 | NA | 109.99 ± 1.96 |
|  | CN<sub>95</sub> | 95 | 225 | 114.14 ± 2.51 | 14.11 ± 0.16 | 28.17 ± 21.78 | NA | 156.43 ± 1.81 |
|  | CN<sub>100</sub> | 100 | 183 | 46.65 ± 1.72 | 9.32 ± 0.19 | 41.58 ± 1.99 | NA | 97.55 ± 2.27 |

**Table 3.**
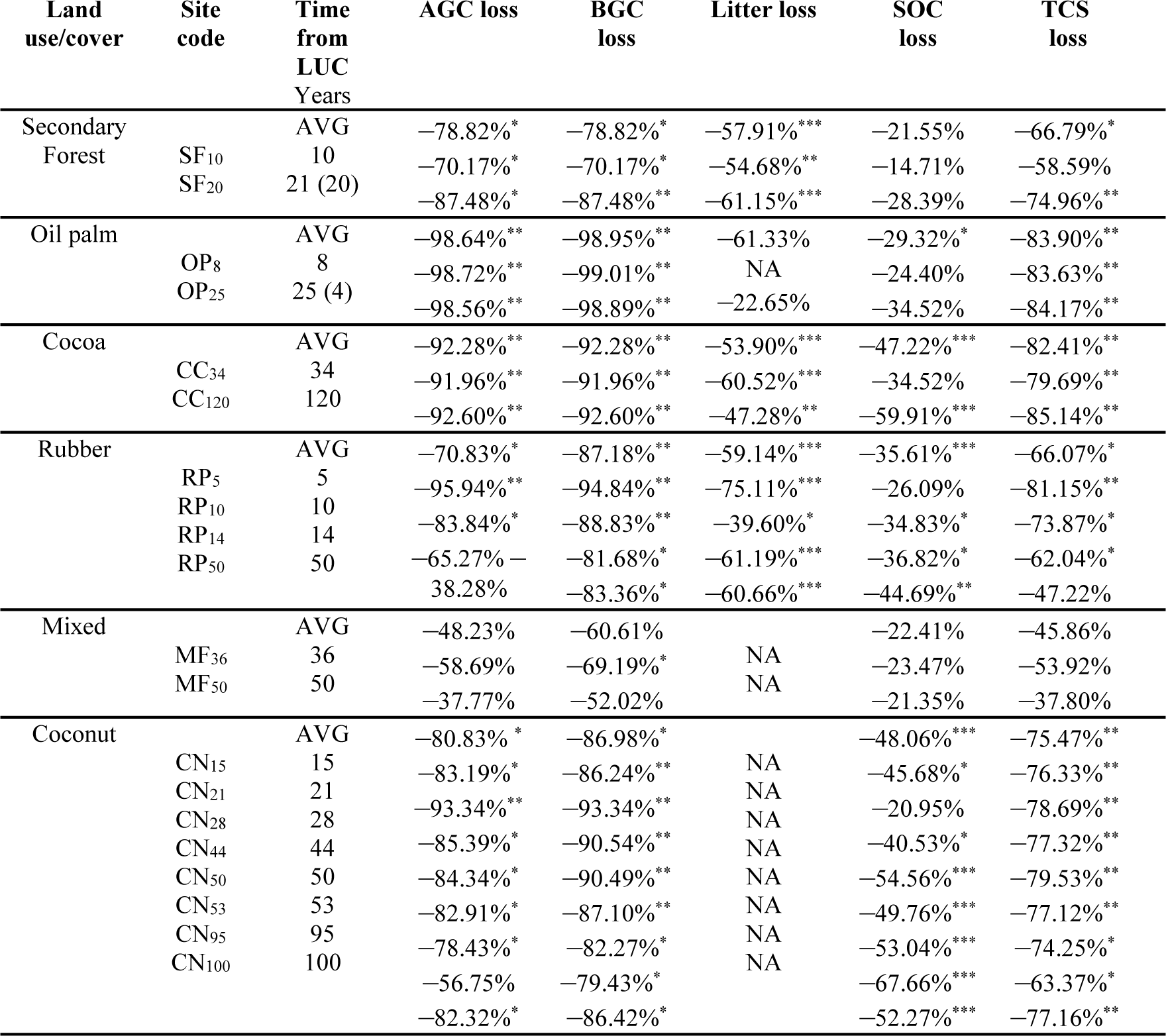
Changes in carbon stock (in %) for different C-pools (AGC, BGC, Litter, SOC) and in TCS considering the primary forest as benchmark reference vs. secondary forest and other tree plantations (positive values represent carbon accumulation and negative values carbon losses compared to the primary forest). The subscript number on the site code indicates the years since deforestation, in parentheses the age of the plantation. AVG are mean losses in % between the primary forest and other land use/cover. ***p-value <0.001, **p-value <0.01, *p-value <0.05. NA = Not Available Data.

#### Below-ground Carbon (BGC)

As for the AGC pool, also BGC results were statistically different for all the land uses compared to the forest with the only exception of mixed plantation (p =0.07)(Figure 3). In this pool, considering the age of the plantations, only the 50-year-old mixed plantation (MP_50_) does not significantly differ from the forest BGC (Figure S1 in Supplementary Material). Overall, forest shows the highest BGC stock (68.24 ±2.20 MgC ha^−1^) among the land uses. Secondary forest, after 10 years from deforestation, accounted for 20.5 ±1.6, while after 21 years, the BGC was 8.69 ±1.4 MgC ha^−1^ with a loss of 70.2 and 87.5%, respectively (Table 3). The estimated BGC annual increment since deforestation was 2 ±0.7 and 0.4 ±0.2 MgC ha^−1^ yr^−1^ in SF_10_ and SF_20_. In the oil palm plantations, BGC was 0.8 ±0.01 (OP_25_, second rotation, 4-year-old plantation) and 0.7 ±0.01 MgC ha^−1^ (OP_8_). BGC loss in the oil palm was respectively 99 and 98.9% while the mean annual increment was 0.2 ±0.1 (OP_25(4)_) and 0.1 ±0.02 (OP_8_) MgC ha^−1^ yr^−1^. Cocoa BGC was estimated for 5.5 ±0.1 (CC_35_) and 5.1 ±0.03 (CC_120_) MgC ha^−1^ with a loss of 92 and 92.6%. The BGC mean annual increment was 0.2 ±0.05 in the 34-year-old plantation and 0.08 ±0.01 MgC ha^−1^ yr^−1^ in the 120-year-old plantation. BGC in rubber plantation ranges from 3.5 ±0.1 (RP_5_) to 12.6 ±0.2 MgC ha^−1^ (RP_14_). The BGB loss varies between 94.8 (RP_5_), to 81.7 (RP_14_), while the estimated mean annual increment was 0.7 ±0.09 (RP_5_), 0.8 ±0.01 (RP_10_), 0.9 ±0.07 (RP_14_) and 0.2 ±0.05 MgC ha^−1^ yr^−1^ (RP_50_), respectively. Mixed plantations had 21.1 ±0.7 (MP_36_) and 32.9 ±1.7 MgC ha^−1^ (MP_50_), with 69.2 and 52% of total loss compared to the forest level with BGC mean annual increment accounting for 0.6 ±0.01 and 0.7 ±0.4 MgC ha^−1^ yr^−1^ on 36-year-old and 50-year-old plantations, respectively. Below-ground carbon stocks in coconut plantations range from 4.6 ±0.02 (CN_22_) to 14.11 ±0.2 (CN_95_) MgC ha^−1^, respectively. The loss for this land use was for all the ages, above 70%, specifically ranging from 79.4 (CN_95_) to 93.3% (CN_21_), respectively, while mean annual BGC increment varies from 0.6 ±0.01 (CN_15_) to 0.1 ±0.05 MgC ha^−1^ yr^−1^(CN_100_)(Table 2 and 3, and Table S2 in Supplementary Material).

**Figure 3.**
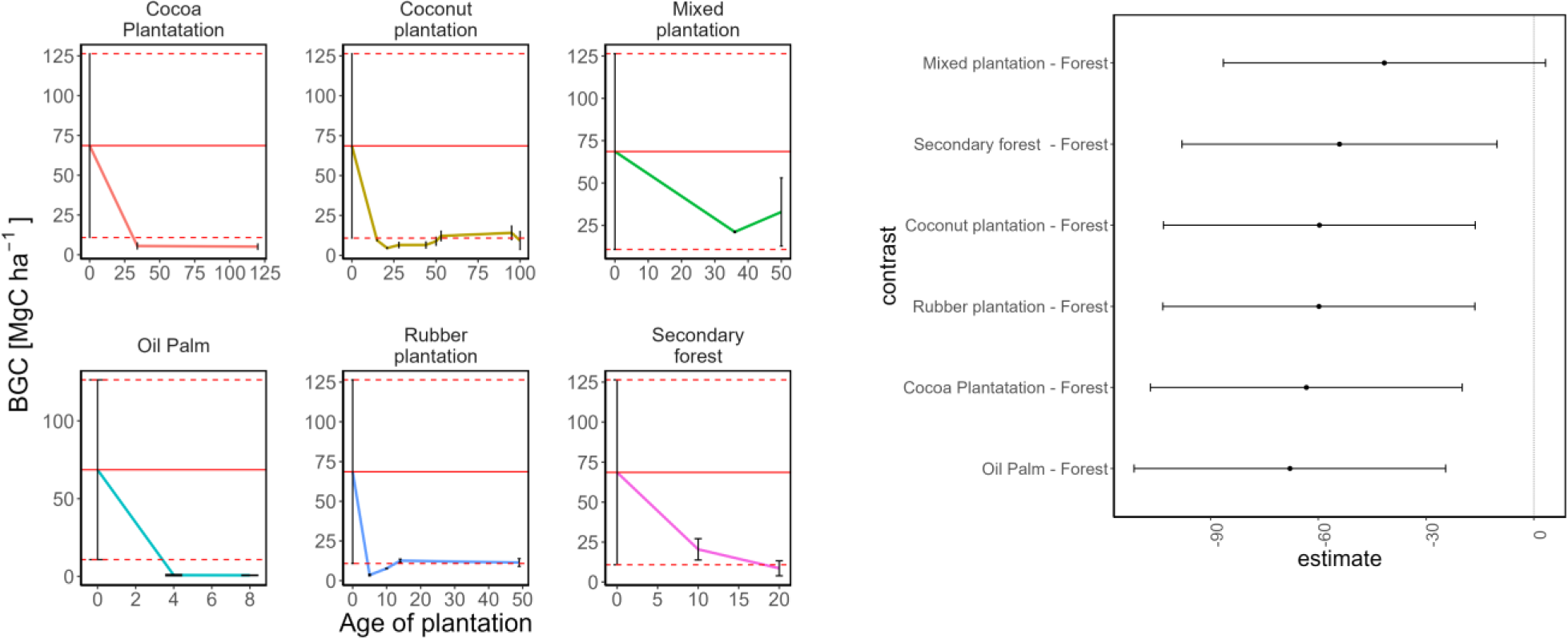
Below-Ground Carbon (BGC in MgC ha^−1^) stocks dynamics in secondary forest and tree plantations (left panel). Statistical significance is represented with Dunnet plot, where confidence interval crossing reference dotted line (0) indicates no statistical difference (p > 0.05)(right panel).

#### Litter carbon

The litter layer was absent in several sites due to common practices of removal, burning and utilization as fodder. Forest litter C-stock was 7.5 ±5.1 MgC ha^−1^, higher than any of the other investigated plantations. No statistically significant differences were found for oil palm plantation (p =0.82)(Figure 4). The litter layer was collected in both the secondary forest sites, accounting for 3.4 ±1.4 in SF_10_ and 2.9 ±0.7 MgC ha^−1^ in SF_21_, with differences compared to the forest of 54.7 and 61.1%, respectively. On oil palm plantations, the litter layer was collected in OP_25_ (4-year-old plantation) since in OP_8_ site was regularly removed for other uses (mainly as fuelwood and fodder). The litter carbon estimated on a leaf number basis results in 5.7 ±3.2 MgC ha^−1^, differing from forest stock by 22.7%. Indeed, cocoa plantations at 35 and 120 years had a litter C-pool accounting respectively for 2.95 ±0.8 and 3.9 ±0.6 MgC ha^−1^, having a percentage difference of 60.5 and 47.3%. Rubber plantations had, in all the sites (5-10-14-50-year-old plantations, respectively), a lower value of carbon litter compared to the forest’s. Litter pools varied from 1.9 ±0.3 in RP_5_ to 4.5 ±1.3 MgC ha^−1^ in RP_10_, while almost identical values are estimated in RP_14_ (2.9 ±0.7 MgC ha^−1^) and RP_50_ (2.9 ±0.8 MgC ha^−1^). This plantation type has a litter C-stock difference, compared to the forest, of about 75.1, 39.6, 61.2 and 60.7% respectively. The litter layer in any of the mixed plantations and coconut sites investigated was regularly removed for other uses (Table 2 and 3, and Table S2 in Supplementary Material).

**Figure 4.**
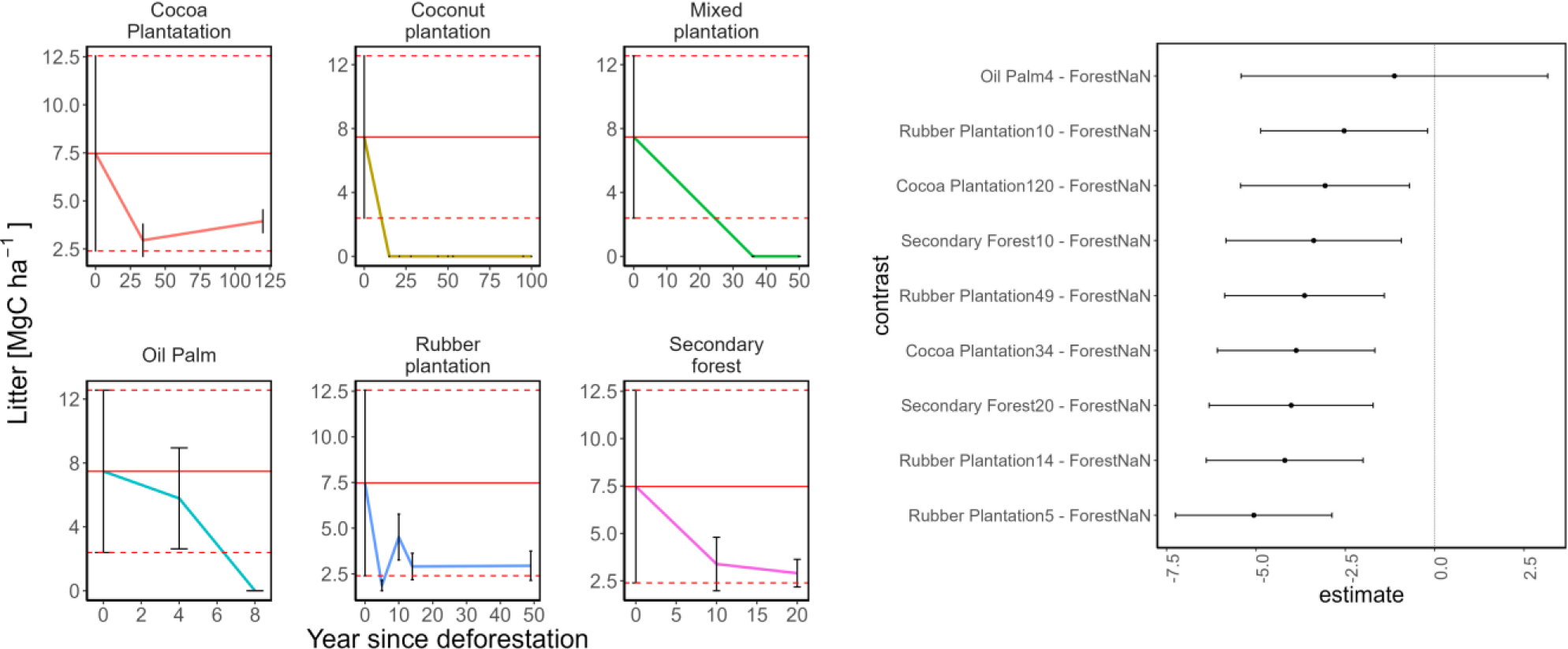
Litter Carbon (in MgC ha^−1^) stocks dynamics in secondary forest and tree plantations (left panel). Statistical significance is represented with Dunnet plot, where confidence interval crossing reference dotted line (0) indicates no statistical difference (p > 0.05)(right panel).

#### Soil carbon

Changes in SOC stocks at 0-30 cm depth, after deforestation, result in a statistically significant decrease (p <0.05) in all cases of conversion into plantations except for mixed plantation (p =0.095) and secondary forest (p =0.115)(Figure 5). Forest SOC in the 0-30 cm layer, accounts for 87.1 MgC ha^−1^ and if considering the age for the comparison with forest SOC stock, no significant differences (p >0.05) are observed with secondary forest (SF_10_ and SF_20_), oil palm (OP_8_ and OP_25(4)_), cocoa (CC_34_), rubber (RP_5_), mixed (MP_36_, MP_50_) and coconut (CN_21_). SOC in secondary forest ranges between 74.3 ±13.9 (SF_10_) and 62.4 ±4.6 MgC ha^−1^(SF_21_) with a total loss of 14.7 and 28.4%, respectively (Table 2, Table 3, Fig. 8). The variation of SOC in the secondary forest is 1.5% (1.3 ±1.4 MgC ha^−1^ yr^−1^) in terms of the mean annual SOC loss and 1.4% (1.2 ±0.2 MgC ha^−1^ yr^−1^) after 10 and 21 years from deforestation. These results are not statistically different from the values of the forest reference level. In oil palm plantations, SOC was 57.3 (OP_25(4)_) and 65.9 MgC ha^−1^ (OP_8_). The decrease in oil palm plantation SOC, after 8 years from deforestation was 21.25 (34.3%) and 29.84 MgC ha^−1^ (24.4%) after 25 years. The mean SOC annual loss was 3% and 1.4% (2.7 ±2 and 1.2 ±1.6 MgC ha^−1^ yr^−1^) in OP_25(4)_ and OP_8_, respectively. SOC in cocoa plantations accounts for 57 ±5.1 and 34.9 ±9.8 MgC ha^−1^ in CC_34_ and CC_120_ respectively. Cocoa plantation, 34 years after deforestation, led to a total SOC decrease of 30.1 MgC ha^−1^ (34.5%) consisting of a mean annual loss of 0.9 ±0.1 MgC ha^−1^ yr^−1^ (1%) while in 120-year-old plantation the total loss accounts for 52.2 MgC ha^−1^ (59.9%) with a mean annual soil loss of 0.43 ±0.08 MgC ha^−1^ yr^−1^ (0.5%). SOC in rubber plantations was 64.4 (RP_5_), 56.8 (RP_10_), 55 (RP_14_) and 48.2 (RP_50_) MgC ha^−1^ but, compared to forest reference level, was observed a SOC loss of 22.7 MgC ha^−1^ (26.1%) in RP_5_, 30.3 (34.8%) in RP_10_, 32.1 (36.8%) in RP_14_ and 38.9 (44.7%) in RP_50_. The mean annual SOC loss was 4.5 ±1.7 (5.2%) in RP_5_, 3 ±0.9 (3.5%) in RP_10_, 2.3 ±0.07 (2.6%) in RP_14_ and 0.78 ±0.05 MgC ha^−1^ yr^−1^ (0.9%) in RP_50_. Mixed plantation had a SOC amounting to 66.7 and 68.5 MgC ha^−1^ after 36 and 50 years from plantation establishment, respectively. SOC decreased of 20.4 MgC ha^−1^ (23.5%) after 36 years with a mean annual loss of 0.57 ±0.3 MgC ha^−1^ yr^−1^(0.7%). After 50 years the total loss accounts for 18.6 MgC ha^−1^ (21.3%) with a mean annual loss of 0.4 ±0.2 MgC ha^−1^ yr^−1^ (0.4%). Differences in SOC between forest and mixed were not statistically significant. The measured SOC in coconut ranges from 28.17 ± 21.78 in CN_95_ to 68.85 ± 7.94 in CN_21_. Comparing to forest level, the depletion in SOC varies from 18.2 (21%) in CN_22_ to 58.9 (67.7%) MgC ha^−1^ in CN_95_. The SOC mean annual loss changes from 2.7 ±0.2 (3%)(CN_15_) to 0.5 ±0.02 (0.5%) (CN_100_)(Table 2 and 3, and Table S2 in Supplementary Material).

**Figure 5.**
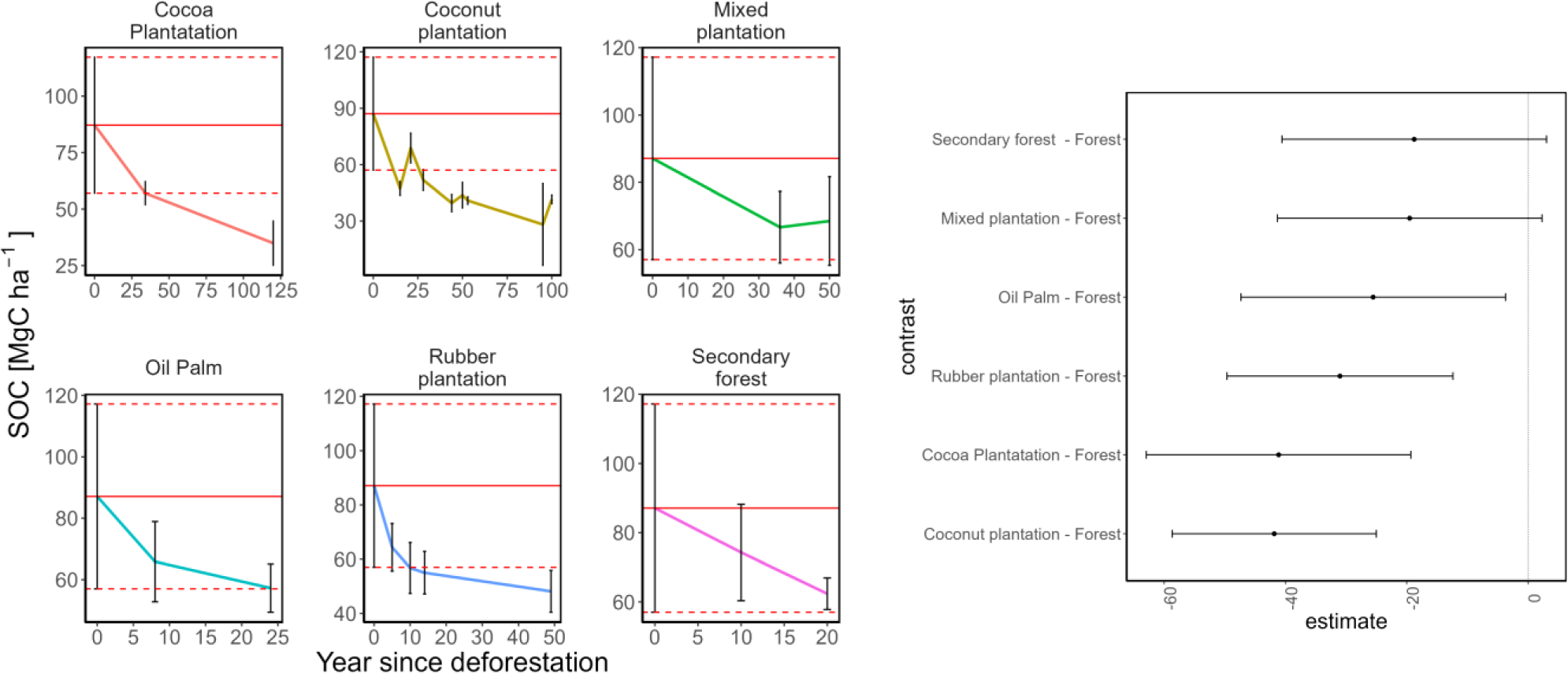
Soil Organic Carbon (SOC in MgC ha^−1^) stocks dynamics in secondary forest and tree plantations (left panel). Statistical significance is represented with Dunnet plot, where confidence interval crossing reference dotted line (0) indicates no statistical difference (p > 0.05)(right panel).

#### Dead standing carbon

The dead wood accounted for 3% of the total biomass in the 120-year-old cocoa plantation and 7% in the 34-year-old cocoa plantation. In coconut plantations, these percentages were 3% (CN_100_) and 6% (CN_53_), while it was 6% for 50-year-old mixed plantations (Table 2, and Table S2 in Supplementary Material).

#### Total carbon

Total carbon differences between forest, secondary forest and tree plantations are all statistically significant (p <0.05), except for mixed plantation (p =0.12)(Figure 6).

**Figure 6.**
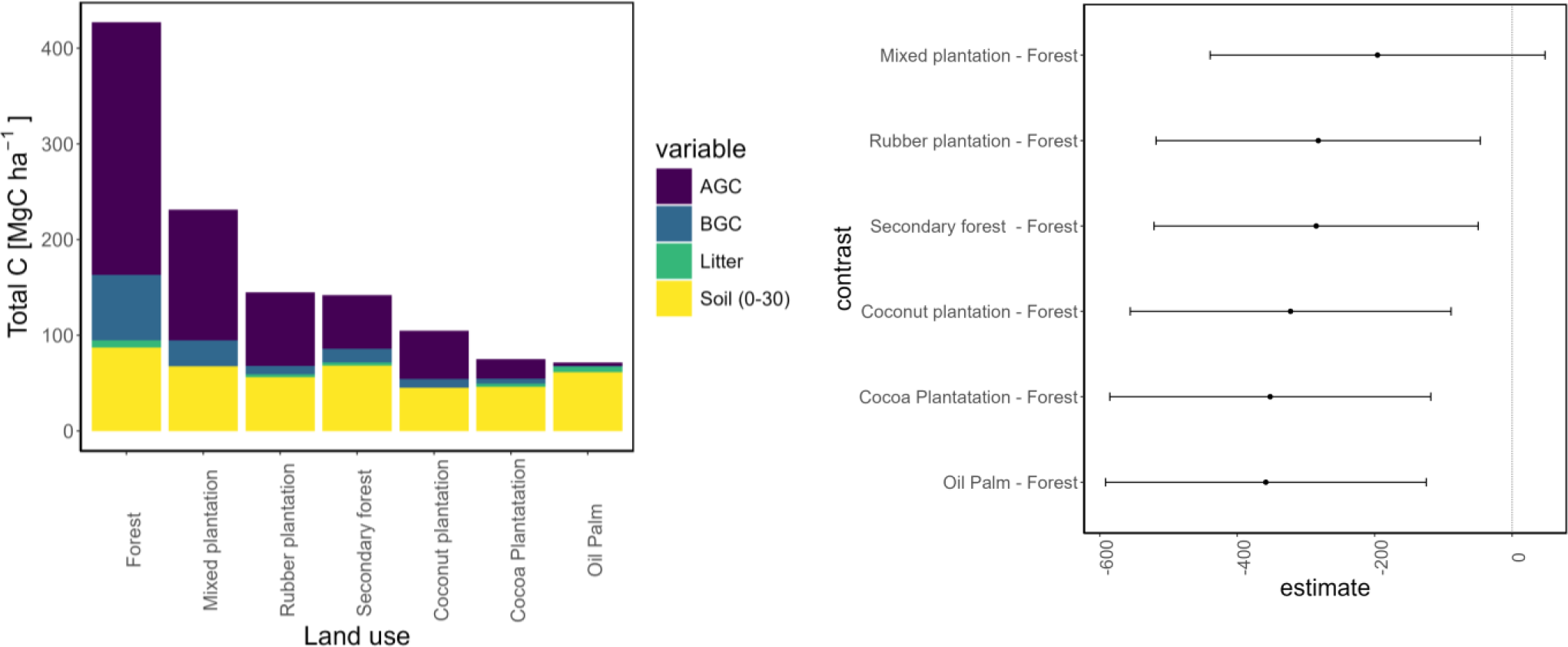
Bar plots represent different Total Carbon Stock (TCS in MgC ha^−1^) in different land uses. Within the same bar, different colours represent different stocks in different C pool (in MgC ha^−1^)(left panel). Statistical significance is represented with Dunnet plot, where confidence interval crossing the reference dotted line (0) indicates no statistical difference (p > 0.05)(right panel).

Considering total carbon (TCS) as the sum of all C-stocks for each pool (but excluding the dead standing wood), forest accounts for 427.1 ±30.6 MgC ha^−1^. This value results higher than any plantation type observed in this study (Figure 7). The estimated TSC in the secondary forest was 176.8 ±14 and 106.9 ±5.6 MgC ha^−1^ with a total C-loss of 250.2 (58.6%) and 320.2 MgC ha^−1^ (75%) in SF_10_ and SF_21_ respectively. For the TCS the estimated loss was 25 ±2 (5.9%) and 15.2 ±1.3 MgC ha^−1^ yr^−1^ (3.6%) after 10 and 21 years since deforestation. Oil palm plantation total carbon was 69.9 ±13.1 (OP_8_) and 67.6 ±7.8 (OP_25(4)_) MgC ha^−1^. These losses, compared to the forest reference level, accounted for 357.2 (83.6%) and 359.5 (84.2%) MgC ha^−1^, respectively, with an annual mean TSC loss of 44.6 ±1.7 (10.5%) and 15 ±3.3 MgC ha^−1^ yr^−1^ (3.5%) after 8 and 25 years from deforestation. Measurement in cocoa showed results of 86.7 ±5.1 and 63.5 ±9.8 MgC ha^−1^, which compared to the forest reference values, represented a loss of 340.4 and 363.6 MgC ha^−1^ in CC_34_ and CC_120_. This means 10 ±0.4 (2.3%) and 3 ±0.1 MgC ha^−1^ yr^−1^ (0.7%) of total mean annual loss, respectively, after 34 and 120 years since forest clearing. Rubber plantation TSC ranges from 80.5 (RP_5_) to 225.4 MgC ha^−1^ (RP_50_) with mean C-losses varying from 346.6 (81.2%) in RP_5_ and 201.7 (47.2%) in RP_50_ Mean annual total C-loss changes from 69.3 ±1.6 (16.2%) in RP_5_ to 4 ±0.9 MgC ha^−1^ yr^−1^ (0.9%) in RP_50_. Mixed plantation presented a TCS of 196.8 ±10.7 and 265.6 ±13 MgC ha^−1^ in MP_36_ and MP_50_ with mean total C-loss of 230.3 (53.9%) and 161.4 MgC ha^−1^ (37.8%) after 36 and 50 years from forest clearing, corresponding to 6.4 ±0.5 (1.5%) and 3.2 ±2.5 MgC ha^−1^ yr^−1^ (0.8%) of mean annual TSC loss. In coconut plantations, TCS ranges from 87.4 ±4.6 (CN_44_) to 156.4 ±21.8 MgC ha^−1^ (CN_95_). The total loss varies from 339.7 (79.5%) in CN_44_ to 270.7 (63.4%) MgC ha^−1^ in CN_95_. The mean annual total carbon loss changes from 21.7 ±0.3 (5.1%) in CN_15_ to 2.8 ±0.6 (0.7%) MgC ha^−1^ yr^−1^ in CN_95_ (Table 2 and 3, and Table S2 in Supplementary Material).

**Figure 7.**
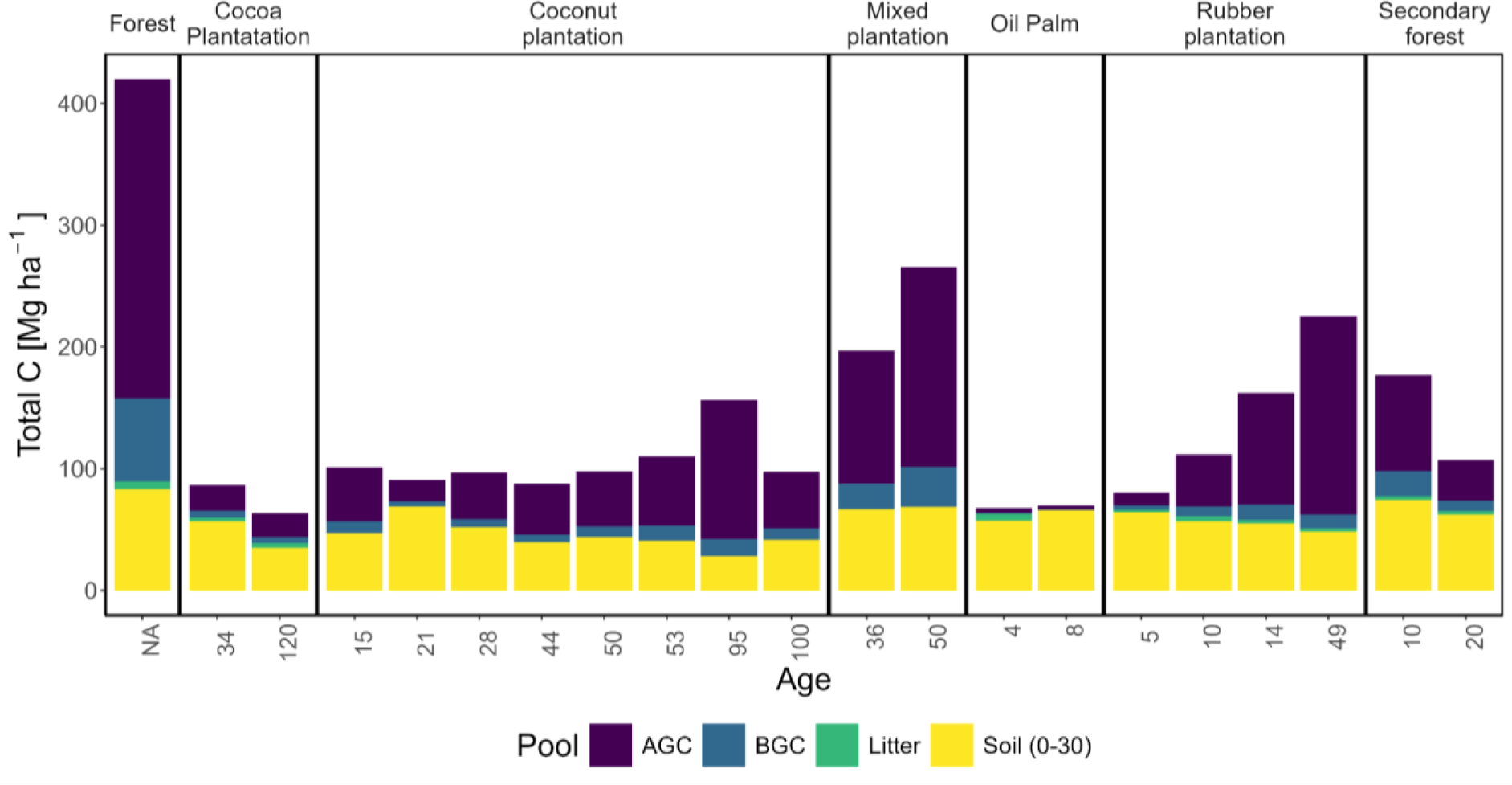
Total Carbon Stocks (TCS = AGC + BGC + SOC + Litter, in MgC ha^−1^) for different land use systems at different stand ages (Age)(AGC = Above Ground Carbon, BGC = Below Ground Carbon, SOC = Soil Organic Carbon).

## 4. Discussions

This work aimed to evaluate the effects of land use change in tropical forest land converted to tree plantations or abandoned to grow as secondary forest. To this aspect, we questioned whether these tropical African ecosystems experience C-stock changes in C-pools at the ecosystem level after deforestation and land use/land cover change and if the range of this variation depended on the pool, the stand age and the type of conversion considered.

### 4.1 Primary forest and mixed plantation lead in C-stock potential

Primary forests are not a direct and immediate source of economic income for local communities especially if compared to tree plantations. In our study, these areas, as well as secondary forests, are considered sacred by community members and only a few activities are allowed (e.g. sporadic hunting and non-timber product harvesting in secondary forests). However, their role at the ecosystem level remains undeniable considering their C-storage potential. Indeed, comparing forest C-stock with secondary forests and tree plantations, we observed that both the total C-stock, and for each single pool, carbon is consistently higher in the primary forest (427.1 ±30.6 vs. 122.6 ±56 MgC ha^−1^ as the mean value of the other TSC’s land uses). Few studies in the literature have been found to consider and describe C-stocks and dynamics in all the C-pools at the same time (i.e. AGC, BGC, Litter and SOC) for identifying the carbon ecosystem potential as we considered here. Our results are, however, within the range observed by Sierra et al. (2007) for a tropical forest in Colombia (299.4 - 467.9 MgC ha^−1^). We found that after deforestation and conversion, only mixed plantations showed no statistically significant differences with forest in terms of total C-stock, and at any of the considered ages of the stand. This appears to align with findings discussed by Hulvey et al. (2013) about the benefits of tree-mixed plantations for their higher C-storage potential compared to monocultures. He et al. (2013) also observed that ecosystem C-storage in mixed plantations is higher than in monoculture in subtropical regions in China, particularly evident considering the soil C-pool. In our study, tree monoculture plantations had similar behaviour to mixed ones with no significative differences compared to forest if not at some specific stand age and for some of the C-pool as in SF_10_(AGB, SOC, TCS), SF_20_(SOC), OP_8_(SOC), RP_5_(SOC), RP_14_(AGC), RP_50_(AGB, TCS), CN_21_(SOC) and CN_95_ (AGB, SOC, TCS).

### 4.2 AGC and SOC are key contributors to total C-stock

In the current study, the estimated reference value of AGC stocks for primary forest is higher than the values observed by Sierra et al. (2007)(247.8 ±40.5 MgC ha^−1^) in Colombia, by Gineste et al. (2008)(from 154.2 to 171.1 MgC ha^−1^) in Ghana, by Lewis et al. (2009) (202 MgC ha^−1^) from 79 plots in tropical African forest and by Adu-Bredu et al. (2010) in Ghana (202.07 MgC ha^−1^) but perfectly in line with values referred to the specific forest-vegetation zone in Ghana by Nyarko et al. (2024)(254 - 278 MgC ha^−1^) and Houssoukpèvi et al. (2022) in Benin (279 ±74 MgC ha^−1^). We estimate that the AGC in primary forest is 61.7% of the total C-stock, followed by SOC (20.4%), BGC (16.1%) and litter carbon (1.7%). Although with different proportions, SOC and AGC are the pools contributing in a larger part to the total ecosystem C-stock in forestland. At the global scale, this is also confirmed by FAO (2020), with 44% of TCS in living biomass, 4% in deadwood, 6% in litter and 45% in soil, respectively. Considering values reported by Sierra et al. (2007), proportions of SOC and AGC were recalculated for a proper comparison, taking into account only SOC in the first 30 cm, resulting in AGB being 44% of the TCS while SOC 38% and BGC 15%. Tree plantations in this study also follow this pattern. It is particularly pronounced the contribution of AGC in older coconut and rubber plantations (73% in CN_95_ and 72.3% in RP_50_) as well as in forest and MP_50_ (both 61.8%). Over 50% of the total C-stock is found in CN_54_ (51.8%), MP_36_ (55.4%) and RP_14_ (56.5%). Rubber proportion of AGC in 14-year-old plantation, perfectly coincides with 56% found by Wauter et al. (2008), for the same stand age in Ghana. Along with AGC, SOC is a prevalent contributor to the total C-stock at the ecosystem level, and this is particularly in younger plantations representing 94.2 and 84.7% of the total C-stock in oil palm plantations (OP_8_ and OP_25(4)_), 80% in RP_5_, 75.7% in CN_22_ and the 65.8% in CC_34_, respectively. These results are similar to those of Houssoukpèvi et al. (2022) who reported SOC contribution to TCS of 63% in tree plantations and young palm SOC contributing for 54% in southern Benin. The higher contribution of SOC to TCS could be attributed to lower AGC due to young stand age and to the original forest SOC still kept after deforestation.

The importance of forest soils at the ecosystem level is widely acknowledged, along with their vulnerability to substantial depletion of SOC upon conversion in agricultural land and tree plantation (Hassink and Whitmore, 1997; Van Noordwijk et al., 2000; Tan et al., 2009; Chiti et al., 2014, 2016; Quezada et al. 2022). In our study, forest SOC values in the first 30 cm lie well within 89.6 ±4.6 MgC ha^−1^ value reported by Chiti et al. (2014) in Ghana and the 84.5 ±40.8 MgC ha^−1^ reported by Ledo et al. (2020) from a global dataset. Higher levels of SOC depletion are found in the secondary forest compared to Don et al. (2011) in the 0-38 cm layer for a 28-year-old secondary forest (12.6 MgC ha^−1^, with a loss of 13% from the previous forest C), perhaps depending on the data derived from 39 different tropical countries. The SOC values we found in the cocoa plantation after 120 years from deforestation were similar to the value of 35.9 ±3.1 MgC ha^−1^ observed in 30-year-old cocoa in Ghana, by Adiyah et al. (2023), at a soil depth of 20-60 cm. Adiyah et al. (2023) presented trends with peculiar directions regarding gain in SOC after an initial decline during the first three years of plantation. We do not have comparable data from young stands but we observed a depletion rate in SOC rate varying from 1 to 0.5% in 34 and 120-year-old plantations respectively, that could not be considered as an increasing trend but probably as a slowing of the rate of loss with plantation because of ageing. The total loss of SOC (0-30 cm) results we found are higher, in absolute values, in older stands but the mean annual loss rate declines with ageing. Indeed, the average annual loss is greater for all plantations during the first years after deforestation with percentages declining from 5.2% (RP_5_) to 0.9% after 50 years (RP_50_). The same trend was estimated in coconut plantations with an annual decrease of 3% in the first 15 years, reaching 0.5% after 100 years. This might suggest that in the long term, the depletion of SOC slows down even though in absolute values of loss, the difference remains greater for older plantations. Comparing even-aged plantations (CN_50_, RP_50_ and MP_50_), SOC depletion is highest in coconut with almost 50% followed by rubber with little less than 45%. Mixed plantations had a no-significant SOC depletion accounting for 21.3%. The absence of statistically significant loss in SOC compared to forest emphasizes the importance of mixed plantations, especially in tropical ecosystems where they are widely adopted by local communities but also because they may represent a win-win solution in terms of livelihoods, food security and, at the same time, for controlling SOC erosion and promoting SOC recovery, overall guaranteeing a higher systemic sustainability.

### 4.3 AGC and BGC accumulation rate is higher in younger stands

Several studies show the prominent effect of age in reducing carbon sequestration capacity and efficiency with ageing (Fonseca et al., 2011; Carbone et al., 2013; Collalti et al., 2020; Luo et al., 2024). Our results show that among all considered land uses/land covers, AGC annual accumulation rate is higher than any other age class in young secondary forest (SF_10_) and young rubber plantation (RP_14_). Secondary forests in humid tropical forest zones are reported to have a greater C-accumulation (due to faster-increasing biomass) during the first 10 years (Fonseca et al., 2011) and then declining with ageing. This is evident in rubber AGC accumulation which we found to be similar to the 5.1 MgC ha^−1^ yr^−1^ reported by Kongsager et al. (2013) for a 12-year-old plantation and to 5.4 MgC ha^−1^ yr^−1^ derived by Wauters et al. (2008) for a 14-year-old rubber plantation (AGC accounting 76.3 MgC ha^−1^) both in western Ghana. With stand ageing, we observed a decline in the accumulation rate for the 50-year-old rubber, nearing the reported values of 3.6 MgC ha^−1^ yr^−1^ by Brahma et al. (2018) in a 40-year-old plantation in India and, similarly, close to the 4.9 MgC ha^−1^ yr^−1^ value reported by Kongsager et al. (2013) in a 40-year-old in Ghana. In our oil palm AGC estimates, values for the 8-year-old are about six times lower than those from Kongsager et al. (2013) for the 7-year-old plantation (21.7 vs. 3.4 MgC ha^−1^). Such a difference may be attributable to a different AGB assessment method highlighting the importance of the method used to estimate the C-pool and the relative uncertainty related to the allometric equations adopted (Vorster et al., 2020). In our case, we did not estimate the AGB through allometric equations but we directly estimated the AGC based on leaves C-content due to the young age of the stands given the absence of stems in such species in the juvenile phase. Another aspect to be considered could be the sites’ peculiarity. Indeed, despite the sites having suitable characteristics for oil palm cultivation, management is undertaken by smallholders, likely with limited resources for ensuring optimal and enhanced plant growth compared to large-scale plantations or, as in the case of Kongsager et al. (2013) study, within an agricultural research station. Considering the annual C-accumulation rate in cocoa plantation we found a reduction in the annual rate from 0.6 for the 34-year-old to 0.2 MgC ha^−1^ yr^−1^ for the 120-year-old plantation. Our values are lower than those reported by Samarriba et al. (2013)(1.3 - 2.6 MgC ha^−1^ yr^−1^) for cocoa plantations in Central America (but under the agroforestry system) and lower than the rate of 3.1 observed by Kongsager et al. (2013) in Ghana and referred to a younger stand (i.e. 21-year-old). Different values could be the result, in the first case, of a more complex and rich stand (agroforestry vs. monoculture) and in the second case could be derived by the younger age in the Kongsager et al. (2013) study. Coconut plantation AGC accumulation rate shows the same declining pattern, as in the other plantations, as the stands grow and reach a maturity stage. Studies from Bhagya et al. (2007) reported 51.14 MgC ha^−1^ for the AGC in 50-year-old stand in India, which resulted in 1.02 MgC ha^−1^ yr^−1^, consistently similar to 0.9 MgC ha^−1^ yr^−1^ from our rate of the same stand age.

A contributing pool to the total ecosystem C-stock is the BGC, which also regulates nutrient cycling and participates in carbon sequestration and climate change mitigation. BGC in cocoa monoculture was estimated at 5.4 MgC ha^−1^ by Borden et al. (2019) in Ghana, which is similar to values ranging from 5.08 to 5.52 MgC ha^−1^ from our study (CC_120_, CC_34_). In the 14-year-old rubber plantation, our estimates of BGC are higher than 7.8 MgC ha^−1^ found by Wauters et al. (2008) in Ghana for the same stand age. We argue that the clone type, which may be different from ours, could affect the results as well as the allometric equations adopted. Overall among the sites, the higher annual accumulation rate of BGC is found in secondary forest. 10 years after forest clearing, the BGC has a comparable value to what was assessed by Brown et al. (2020) for a 44-year-old secondary forest (20.5 vs. 20.7 MgC ha^−1^) in Ghana. All the rest of the sites had shown annual accumulation rates lower than 1 MgC ha^−1^, implying a less predominant role in the TCS balance. No significant differences were found in MP_50_ compared to the reference level of BGC in the forest, following the same outcomes as for AGC.

## 5. Conclusions

Deforestation, land use changes and their impact on carbon stock dynamics in various ecosystems over time play a pivotal role in both development policies and natural resources conservation strategies. Particularly in tropical ecosystems, anthropogenic pressure is leading to a progressive reduction of forested areas which are replaced by tree monoculture, pastures, crops, mixed and agroforestry plantations, from both smallholders and big enterprises. Our findings highlight the potentiality of C-sequestration of some of the main and more common land uses in tropical ecosystems. Although the primary forest ecosystem has the highest amount of carbon in any of the assessed pools, taking into consideration the more relevant pools in terms of contribution to the total ecosystem C-stock (AGC and SOC) no significative differences were found in tree mixed plantation and secondary forest. This highlights the crucial role of local smallholders in preserving and restoring C-stock ecosystems, alongside the key role of local government in protecting and enhancing forest resources. Our work also highlights the need to deepen the knowledge about the role of secondary forests as carbon reservoirs, particularly in the tropics, where they are largely understudied. The outcomes from the rest of the land uses are related to the age of the plantation, underlining that the SOC component, higher in younger stands, may represent residue of the previous forest cover asset which decreases over time once a plantation is established. Conversely, the AGC grows, with the plantation ageing after deforestation (e.g. in rubber and mixed plantations) with a declining rate as the stand ages. Our work also highlights the growing need for carbon estimates that account for and consider all the C-pools within the ecosystem and not only a fraction of them, this to better quantify the overall C-stock and sequestration capacity, given that, as we demonstrated here, they can vary (much) between the different carbon pools.

Tree crop plantations, particularly mixed stands, may be beneficial for smallholder livelihood, food security as well as carbon sequestration potential. However, land use changes need to be properly planned and tree plantations possibly established on lands with low carbon content such as degraded or agricultural lands. While tree plantations may not match primary forests in climate change mitigation, some could efficiently meet both the local community livelihood needs and enhance carbon stock if not established by clearing primary forests. A full comprehension of the ecosystem C-sequestration potential and C-stock dynamics after deforestation is a key point in developing sustainable mitigation strategies and forest protection planning. This could be achieved with combined efforts in assessing the complex and intricate environmental and socio-economic interactions within deforestation-risk lands, in a comprehensive view for effective and long-term sustainable rural development.

## Acknowledgements

Fieldwork was funded by grants from Africa-GHG: 247349 under the European Union’s Seventh Framework Programme (FP7/2007-2013).

E.G. and A.C. also acknowledge funding by the project OptForEU Horizon Europe research and innovation programme under grant agreement No. 101060554. E.V. and A.C. also acknowledge funding from PRIN 2020 (cod 2020E52THS) - Research Projects of National Relevance funded by the Italian Ministry of University and Research entitled: “Multi-scale observations to predict Forest response to pollution and climate change” (MULTIFOR, project number 2020E52THS). T.C. and A.C. also acknowledge the project funded under the National Recovery and Resilience Plan (NRRP), Mission 4 Component 2 Investment 1.4 - Call for tender No. 3138 of 16 December 2021, rectified by Decree n.3175 of 18 December 2021 of Italian Ministry of University and Research funded by the European Union – NextGenerationEU under award Number: Project code CN_00000033, Concession Decree No. 1034 of 17 June 2022 adopted by the Italian Ministry of University and Research, CUP B83C22002930006, Project title “National Biodiversity Future Centre - NBFC”.

We would like to gratefully acknowledge the Wildlife Division of the Forestry Commission in Ghana, Ankasa Park managers Kareem A. F. and Balangtaa C. and all the park rangers for their unwavering helpfulness. Special thanks go to the field team Mensa J.J., Cudjoe E. and Cobienna J. who helped with data collection on the field. Gratitude and great appreciation are expressed to the local communities and traditional authorities, for their warm welcome and trust, which made our work possible.

## Author contributions

**Elisa Grieco:** Conceptualization, Data collection and data curation, Formal analysis, Investigation, Writing - original draft. **Elia Vangi:** Formal analysis, Investigation, Writing - original draft. **Tommaso Chiti:** Data collection and data curation, Formal analysis, Investigation, Writing - original draft. **Alessio Collalti:** Conceptualization, Formal analysis, Investigation, Writing - original draft

