## Supplementary Material for "Impacts of Deforestation and Land Use/Land Cover Change on Carbon Stock and Dynamics in the Jomoro District, Ghana"

**Grieco et al.**

### Supplementary material

**Table S1**. Selected sites in Jomoro district (in parentheses the age of the plantation)

| Land use/cover | Site code | Years from def. | Locations | | |
|---|---|---|---|---|---|
| Forest | ANKF, | n.a. | 05.130115N - 002.385363W | 05.130024N - 002.385080W | 05.125766N - 002.385014W |
| | CTF, | | 05.192981N - 002.453037W | 05.192844N - 002.452989W | 05.192896N - 002.453102W |
| | NANAF | | 05.06069N - 002.53348 W | | |
| Secondary Forest | $SF_{10}$ | 10 | 05.20214N - 002.64112W | 05. 20208N - 002.64081W | 05. 20241N - 002.64062W |
| | $SF_{20}$ | 21 (20) | 05.12743N - 002.51182W | 05.12727N - 002.51162W | 05.12702N - 002.51185W |
| Oil palm | $OP_{8}$ | 8 | 05.132998N - 002.432988W | 05.133088N - 002.432858W | 05.133088N - 002.432858W |
| | $OP_{25}$ | 25 (4) | 05.20968N - 002.64966W | 05.20983N - 002.64912W | 05.21004N - 002.64899W |
| Cocoa | $CC_{120}$ | 120 | 05.192277N - 002.461279W | 05.192277N - 002.461279W | 05.18361N - 002.71775W |
| | $CC_{34}$ | 34 | 05.18156N - 002.71925W | 05.18376N - 002.71875W | 05.18361N - 002.71775W |
| Rubber | $RP_{5}$ | 5 | 05.14492N - 002.51010W | 05.14451N - 002.51089W | 05.14578N - 002.51013W |
| | $RP_{10}$ | 10 | 05.124770 N - 002.415371W | 05.124495N - 002.415419W | 05.124495N - 002.415617W |
| | $RP_{14}$ | 14 | 05.06170N - 002.53896W | 05.06012N - 002.53811W | 05.06111N - 002.53830W |
| | $RP_{50}$ | 50 | 05.10176N - 002.59658W | 05.10344N - 002.59596W | 05.10446N - 002.59692W |
| Mixed | $MP_{36}$ | 36 | 05.13937 N - 002.50424 W | 05.13931 N - 002.50357 W | 05. 13884 N - 002.50316 W |
| | $MP_{50}$ | 50 | 05.17448 N - 002.65448 W | 05.17475 N - 002.65513 W | 05. 17444 N - 002.65404 W |
| Coconut | $CN_{15}$ | 15 | 05.042529N - 002.395288W | 05.042624N - 002.395099W | 05.042599N - 002.394805W |
| | $CN_{21}$ | 21 | 05.20117N - 002.64137W | 05.20037N - 002.64102W | 05.20063N - 002.64064W |
| | $CN_{28}$ | 28 | 05.102258N - 002.390405W | 05.101864N - 002.390306W | 05.102288N - 002.390768W |
| | $CN_{44}$ | 44 | 05.15926N - 002.62520W | 05.15825N - 002.62556W | 05.15720N - 02.62694W |
| | $CN_{50}$ | 50 | 05.100052N - 002.375682W | 05.10020N - 002.380050W | 05.095658N - 002.375749W |
| | $CN_{53}$ | 53 | 04.99522N - 002.60818W | 04.99642N - 002.60809W | 04.99512N - 002.60907W |
| | $CN_{95}$ | 95 | 05.005614N - 002.432568W | 05.005353N - 002.432906W | 05.005292N - 002.432211W |
| | $CN_{100}$ | 100 | 05.040517N - 002.362435W | 05.040386N - 002.362628W | 05.040176N -002.362587W |

**Table S2**. Rates of mean annual carbon stock changes (in MgC ha$^{-1}$ yr$^{-1}$)(positive values represent carbon accumulation while negative values represent carbon losses) in different pools (AGC, BGC, Litter, SOC) and in TCS for the secondary forest and tree plantations in comparison with the primary forest considered as the benchmark reference. The subscript number on the site code indicates the years since deforestation, in parentheses the age of the plantation. '±' denotes one standard deviation. ***p-value <0.001, **p-value <0.01, *p-value <0.05. NA = Not Available Data.

| **Land use/cover** | **Site code** | **Time from LUC** Years | **AGC** | **BGC** | **Litter** | **SOC** | **TCS** |
|---|---|---|---|---|---|---|---|
| Secondary Forest | $SF_{10}$ | 10 | 7.9 ± 2.6 (10%) | 2.0 ± 0.7 (10%) | 0.41 ± 0.1 (12%) | –1.3 ± 1.4 (1.5%) | –25 ± 2.0 (5.9%) |
| | $SF_{20}$ | 21 (20) | 1.7 ± 1.3 (5%) | 0.4 ± 0.2 (5%) | 0.23 ± 0.03 (8%) | –1.2± 0.2 (1.4%) | –15.2 ± 1.3 (3.6%) |
| Oil palm | $OP_{8}$ | 8 | 0.4 ± 0.1 (12.5%) | 0.1 ± 0.02 (12.5%) | NA | –2.7 ± 1.6 (3%) | –44.6 ± 1.7 (10.5%) |
| | $OP_{25}$ | 25 (4) | 1.0 ± 0.5 (25%) | 0.2 ± 0.1 (25%) | 0.42 ± 0.8 (7%) | –1.2 ± 1.9 (1.4%) | –15 ± 3.3 (3.5%) |
| Cocoa | $CC_{34}$ | 34 | 0.6 ± 0.2 (2.9%) | 0.2 ± 0.05 (2.9%) | 0.13 ± 0.02 (5%) | –0.9 ± 0.15 (1%) | –10 ± 0.4 (2.3%) |
| | $CC_{120}$ | 120 | 0.2 ± 0.05 (0.8%) | 0.04 ± 0.01 (0.8%) | 0.03 ± 0.005 (1%) | –0.4 ± 0.08 (0.5%) | –3.0 ± 0.1 (0.7%) |
| Rubber | $RP_{5}$ | 5 | 2.1 ± 0.4 (20%) | 0.7 ± 0.09 (20%) | 1.12 ± 0.06 (60%) | –4.5 ± 1.7 (5.2%) | –69.3 ± 1.6 (16.2%) |
| | $RP_{10}$ | 10 | 4.3 ± 0.5 (10%) | 0.8 ± 0.01 (10%) | 0.30 ± 0.1 (7%) | –3.0 ± 0.9 (3.5%) | –31.5 ± 0.6 (7.4%) |
| | $RP_{14}$ | 14 | 6.5 ± 1.4 (7.1%) | 0.9 ± 0.07 (7.1%) | 0.33 ± 0.05 (11%) | –2.3 ± 0.5 (2.6%) | –18.9 ± 0.9 (4.4%) |
| | $RP_{50}$ | 50 | 3.3 ± 0.7 (2%) | 0.2 ± 0.05 (2%) | 0.09 ± 0.2 (3%) | –0.8 ± 0.15 (1%) | –4.0 ± 0.9 (0.9%) |
| Mixed | $MF_{36}$ | 36 | 3.0 ± 0.5 (2.8%) | 0.6 ± 0.01 (2.8%) | NA | –0.6 ± 0.3 (0.7%) | –6.4 ± 0.5 (1.5%) |
| | $MF_{50}$ | 50 | 3.3 ± 2.1 (2%) | 0.7 ± 0.4 (2%) | NA | –0.4 ± 0.3 (0.4%) | –3.2 ± 2.5 (0.8%) |
| Coconut | $CN_{15}$ | 15 | 3.0 ± 0.08 (6.7%) | 0.6 ± 0.01 (6.7%) | NA | –2.6 ± 0.2 (3%) | –21.7 ± 0.3 (5.1%) |
| | $CN_{21}$ | 21 | 0.8 ± 0.07 (4.5%) | 0.2 ± 0.01 (4.5%) | NA | –0.8 ± 0.4 (1%) | –15.3 ± 0.3 (3.6%) |
| | $CN_{28}$ | 28 | 1.4 ± 0.4 (3.6%) | 0.2 ± 0.06 (3.6%) | NA | –1.3 ± 0.2 (1.4%) | –11.8 ± 0.4 (2.8%) |
| | $CN_{44}$ | 44 | 0.9 ± 0.2 (2.3%) | 0.1 ± 0.04 (2.3%) | NA | –1.1 ± 0.1 (1.2%) | –7.7 ± 0.1 (1.8%) |
| | $CN_{50}$ | 50 | 0.9 ± 0.2 (2%) | 0.2 ± 0.05 (2%) | NA | –0.9 ± 0.1 (1%) | –6.6 ± 0.4 (1.5%) |
| | $CN_{53}$ | 53 | 1.1 ± 0.09 (1.9%) | 0.2 ± 0.05 (1.9%) | NA | –0.9 ± 0.04 (1%) | –6.0± 0.1 (1.4%) |
| | $CN_{95}$ | 95 | 1.2 ± 0.5 (1.1%) | 0.1 ± 0.04 (1.1%) | NA | –0.6 ± 0.2 (0.7%) | –2.8 ± 0.6 (0.7%) |
| | $CN_{100}$ | 100 | 0.5 ± 0.2 (1.0%) | 0.1 ± 0.05 (1.0%) | NA | –0.6 ± 0.02 (0.5%) | –3.3± 0.3 (0.8%) |

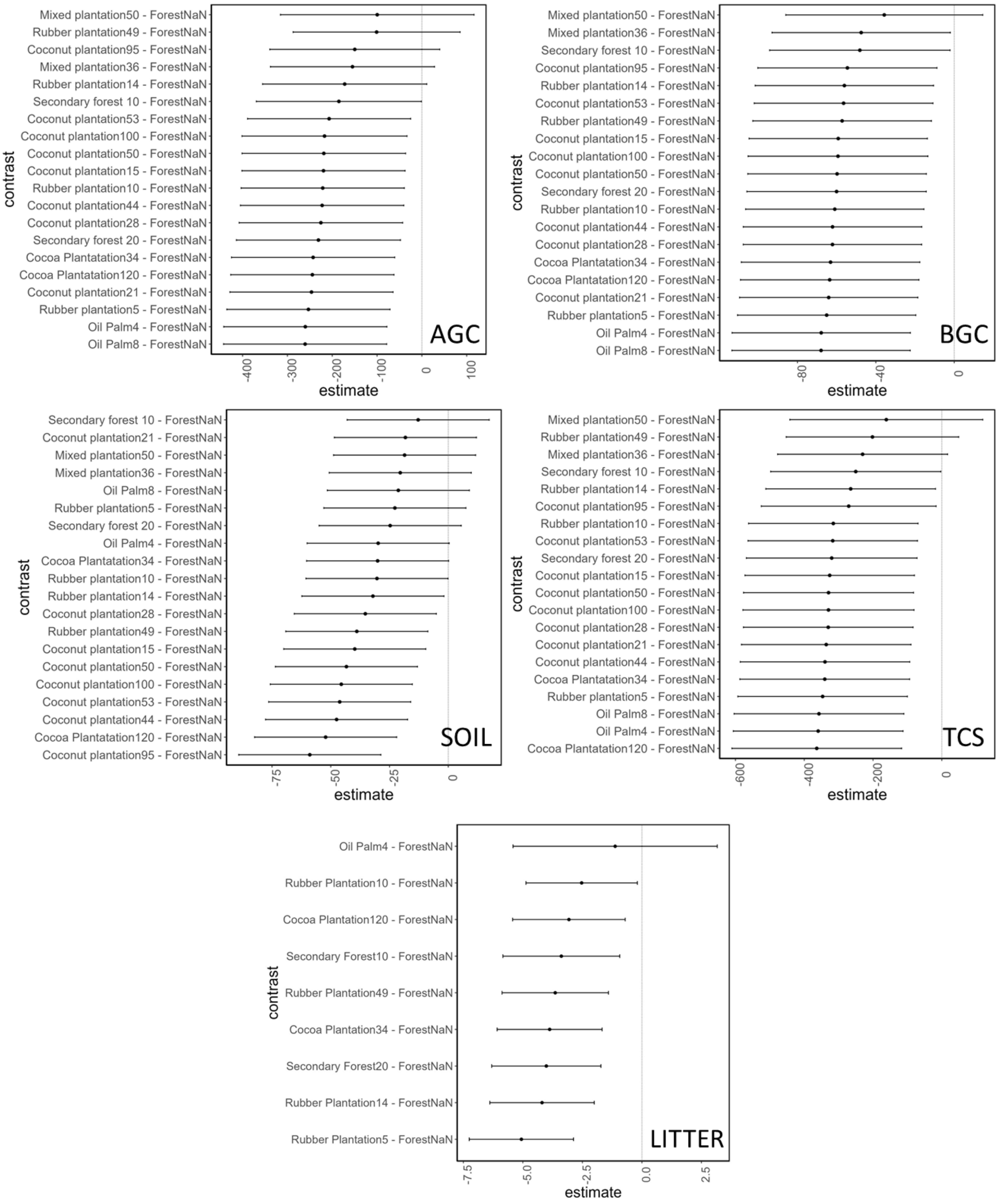


**Figure S1**. Statistical significance of different stand ages compared to the primary forest, based on different C pools, represented with Dunnet plot, where confidence interval crossing reference dotted line (0) indicates no statistical difference ($p > 0.05$).
